# Tumour suppressor WT1 regulates the let-7-*Igf1r* axis in kidney mesenchyme

**DOI:** 10.1101/822973

**Authors:** Ruthrothaselvi Bharathavikru, Joan Slight, Stuart Aitken, Giulia Petrovich, Jocelyn Charlton, Viktoriya Stancheva, Abdelkader Essafi, Kathy Pritchard-Jones, Nicholas D Hastie

**Author notes:** Department of Biological sciences, Indian Institute of Science Education and Research, Berhampur, Ganjam, Odisha 760010 (Ruthrothaselvi Bharathavikru).

## Abstract

Wilms’ tumour 1 (WT1) is a transcription factor and a tumour suppressor, essential for the development and homeostasis of multiple tissues derived from the intermediate and lateral plate mesoderm. Germline *WT1* mutations result in the eponymous paediatric kidney cancer, genitourinary anomalies and in some cases congenital diaphragmatic hernia One common feature in Wilms’ Tumours (WT), is upregulation of IGF2 through genetic and/or epigenetic mechanisms. Recent studies have identified both somatic and germline mutations in microRNA processing genes (MIRPG) in WT. Whether these different epigenetic and genetic causes converge on common targets and the mechanisms by which they act are still unclear. WT1 is involved in RNA binding and regulates the RNA stability of important developmental genes. We now show that WT1 interacts with let-7 family of microRNAs, and the absence of WT1 results in reduced levels of mature microRNA in cell lines and kidney mesenchyme. As a consequence, let-7 targets, including Igf1 receptor (*Igf1r*), are upregulated in the absence of *Wt1*, thus confirming the presence of a WT1-let7-*Igf1r* axis. These findings suggest a possible mechanism by which *WT1* mutations lead to WT, and reinforce the idea that the perturbation of the microRNA and IGF signalling pathways are important contributing factors in the aetiology of WT.

## Introduction

Wilms’ Tumour (WT) is a paediatric kidney disease that may arise from germline mutations in the tumour suppressor genes *WT1* or *WTX*. The incidence of WT is approximately one in every 10,000 (Charlton and Pritchard-Jones, 2016). Germline mutations of *WT1* also lead to other genitourinary abnormalities, and, in some rare cases, congenital diaphragmatic hernia (Hastie, 2017). Some of the overgrowth syndromes, such as Beckwith–Wiedemann Syndrome (BWS) and Perlman Syndrome, also have an increased risk of Wilms’ Tumour. The mechanisms contributing to WT are complex. For example, both *WT1* mutations and *IGF2* upregulation (brought about by genetic and/or epigenetic changes), have now been shown to be present in WT as well as BWS patients. This is also true in the mouse model, where *Wt1* deletion along with activation of Igf2 signalling leads to WT (Hu et al., 2011). Other categories of mutations include stabilization of β catenin and *p53* mutations (especially in some anaplastic tumours) (Bardeesy et al., 1994).

Recent studies with patient samples have used targeted exome sequencing to categorize additional mutations in WT. Apart from the genes that are involved in kidney development, a new category of mutations in the microRNA processing pathway (Wegert et al., 2015, Gadd et al., 2017, Ludwig et al., 2016) was identified. Thus, the driver mutations can now be divided into two major categories; nephron differentiation (*WT1, SIX2, METT*), and the microRNA processing gene (MIRPG) pathway (*DICER, DIS3L2, DROSHA, DGCR8*). The microRNA processing pathway is a highly co-ordinated regulatory programme that regulates RNA levels, thereby affecting protein levels as well. The microRNA pathway consists of the following steps: i) transcription of the primary microRNA (pri-miRNA) from the genome by a Pol-II driven mechanism, ii) followed by the action of the microprocessor complex, composed of DGCR8 and DROSHA, which recognises the stem loop structure and the sequence of the pri-miRNA, which is then cleaved to form the premature microRNA (pre-miRNA), iii) The pre-miRNA is then exported to the cytoplasm by the exportin 5 complex, iv) where the double-stranded pre-miRNA is then recognized by DICER, which along with other proteins, further processes it and forms mature microRNA. The mature microRNA consists of two strands, the guide RNA and the passenger RNA, v) The guide RNA is then loaded onto the RNA induced silencing complex (RISC) complex, that comprises Argonaute (AGO 1/2). The RISC is then capable of silencing gene expression by different mechanisms that include deadenylation, translational repression and degradation mainly by targeting the 3’UTR of the mRNAs (Winter et al., 2009).

The *MIRPG* mutations identified in WT are mostly somatic, except those in *DICER*, which in some cases were also germline (Palculict et al., 2016). The microRNA pathway has also been indirectly linked to WT. Recent studies have shown that the MIRPG-mutation associated with WTs also converge on the IGF signalling pathway through transcriptional mechanisms (Hunter et al., 2018). The Perlman overgrowth syndrome, characterized by an increased susceptibility to WT, arises because of mutations in the exoribonuclease *DIS3L2*, which is involved in degradation of oligouridylated transcripts, including microRNA let-7 (Astuti et al., 2012, Chang et al., 2013). One of the recent mouse models of WT was created by overexpressing LIN28 in the kidney mesenchyme where WT1 is usually expressed (Urbach et al., 2014). The RNA binding protein LIN28 is involved in the homeostasis of the microRNA let-7 pathway by binding to let-7 microRNAs and preventing their translational repression (Tsialikas and Romer-Seibert, 2015, Viswanathan et al., 2008). Although a global reduction in microRNA levels was observed in the LIN28 mouse model study, the introduction of let-7g resulted in tumour regression, suggesting the let-7-LIN28 microRNA pathway to be the one of the underlying cause of WT manifestation and tumour pathogenesis.

Wilms Tumour 1 (WT1) protein is one of the major tumour suppressor proteins that is associated with WT. The transcription factor role of WT1 has been considered to be responsible for the cellular changes observed in WT. However, we have recently shown that WT1 is also a regulator of RNA stability and that its role in posttranscriptional regulation is equally important in development (Bharathavikru et al., 2017). We had also observed in this study that WT1 interacts with microRNAs. We decided to investigate if WT1 played a role in the microRNA processing pathway, especially in the let-7 microRNA pathway, since the let7-LIN28 pathway has been previously shown to be associated with WT.

WT1 interaction with microRNAs is enriched in the cytoplasmic compartment suggesting that the interaction occurs after the microprocessor activity. Detailed analysis of microRNA expression shows that, in the absence of WT1, there is a defect in the microRNA processing step, which can be regained by a transient induction of WT1. The most significant category of microRNAs that responds to the levels of WT1 is the let-7 family. WT1 interacts with the microRNA processing pathway proteins and thus modulates the microRNA processing pathway. The microRNA processing defect is also observed in kidney mesenchyme deleted for *Wt1* using a *Nestin-cre Wt1* conditional mouse model. In WTs, there is a decrease in the mature microRNA let-7 levels in samples with *WT1* mutation or loss. Further analysis of let-7 targets shows a significant upregulation of *Igf1r* that impinges on the IGF signalling pathway. All these observations suggest that the dysregulation of the microRNA let-7 pathway and the downstream *Igf1r* modulation is one of the contributing causes of WT.

## Results

### Tumour suppressor WT1 interacts with microRNAs

WT1 regulates developmental targets through transcriptional as well as posttranscriptional mechanisms, influencing mRNA stability through direct interactions with the 3’ UTR (Bharathavikru et al., 2017). The global analysis of WT1-interacting RNA (**Fig.S1A**), by FLASH (<u>F</u>orma<u>L</u>dehyde-<u>A</u>ssisted crosslinking and <u>S</u>equencing of <u>H</u>ybrids) and RNA immunoprecipitation (RIP), was done in mesonephric M15 cells (Bharathavikru et al., 2017). This showed different noncoding and small RNA subtypes, of which microRNA was one of the common enriched categories (**Fig. 1A, S1B**). Due to the technical differences of the two RNA pulldown techniques, the proportion of the subtypes identified by each technique differ. The comparison of WT1-interacting microRNA to the input control identified by FLASH is depicted in **Fig. 1B**, which shows a characteristic bimodal pattern of enrichment. The interaction with microRNAs was confirmed by northern blotting using probes specific for the mature microRNAs let-7c, let-7g and U6 (**Fig. 1C, S1C).** The microRNA-processing pathway has both nuclear and cytoplasmic components. The nuclear compartment is the site for the pri-microRNA to pre-microRNA processing. The pre-microRNA is then transported to the cytoplasm, where the pre-microRNA is processed to mature microRNA (**Fig.1D**). WT1 is known to shuttle between the nucleus and cytoplasm (Niksic et al., 2004); and hence, we performed subcellular fractionation (**Fig.S1D**) followed by WT1 RIP to identify the compartment where WT1-microRNA interaction occurs. WT1-interacting microRNAs were analysed, using primers specific for the pre-processed and processed microRNAs as depicted in **Fig. S1A**. Both the premature and the mature microRNA let-7c were both found to interact with WT1, and the maximum enrichment of WT1-interacting microRNAs was identified in the cytoplasmic compartment (nearly 100 fold more than the nuclear compartment) (**Fig. 1D**). This indicates that the WT1 microRNA-interaction occurs from the DICER node of the microRNA processing pathway.

**Figure 1:**
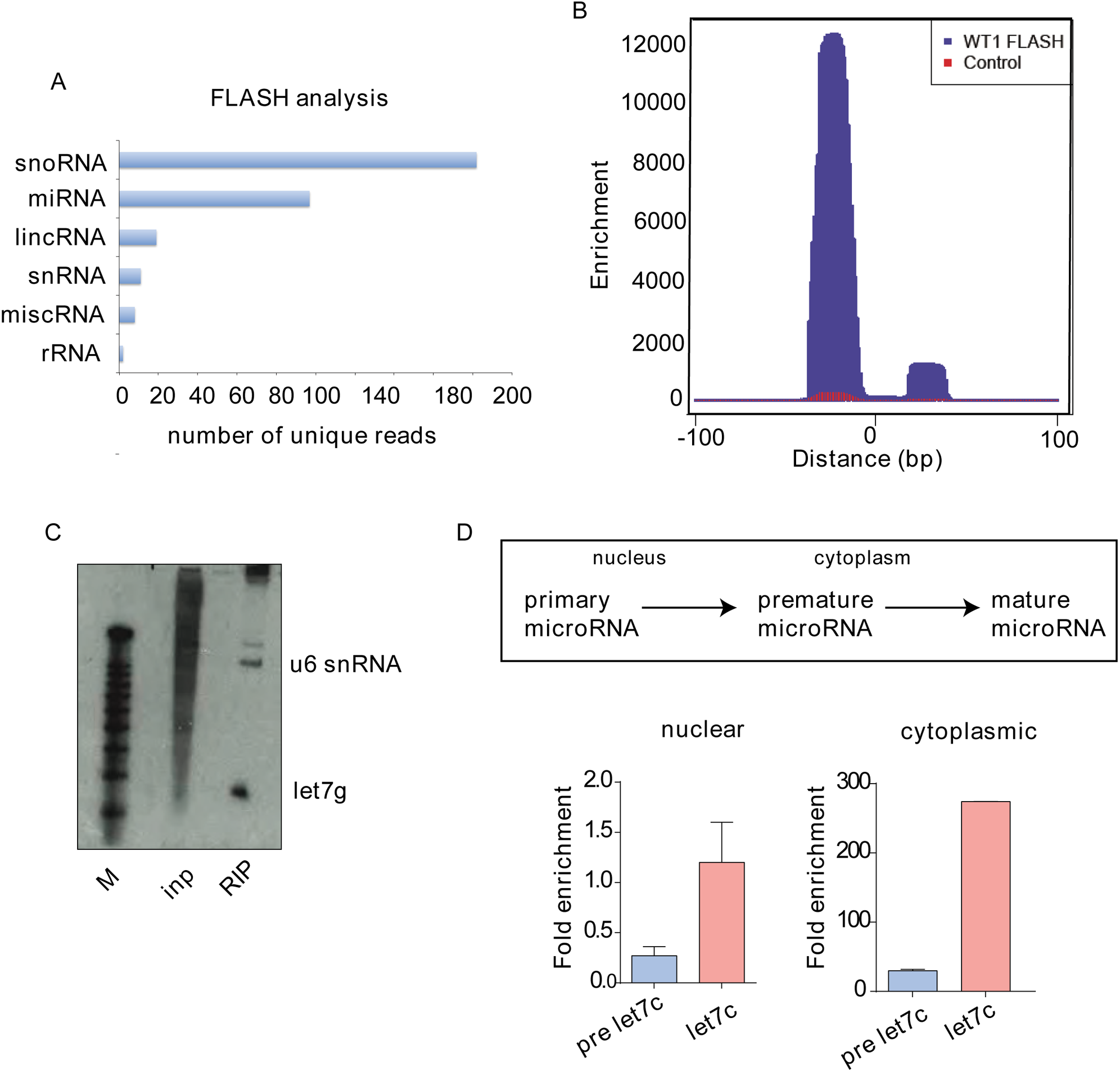
Tumour suppressor WT1 interacts with microRNAs. A) Graphical representation of the percentage of WT1 interacting noncoding biotypes; lincRNA, miRNA, rRNA, snRNA, snoRNA, misc RNA in FLASH (<u>F</u>orma<u>L</u>dehyde-<u>A</u>ssisted crosslinking and <u>S</u>equencing of <u>H</u>ybrids) analysis of mesonephric cell line, M15. B) Read pile-up plot of microRNAs in M15 WT1 FLASH analysis compared to input control. The x-axis represents the 200 base pair window, centered in the middle of each miRNA. C) Northern blotting analysis of WT1 interacting let-7g microRNA compared to input sample. U6 snRNA is also detected as an interacting noncoding RNA in the same sample. D) WT1 interacts preferentially with pre-miRNA let-7c and mature miRNA let-7c, in the cytoplasmic compartment as identified by WT1 RNA pulldowns from subcellular compartments. The different subcellular location of the primary, pre and mature microRNA is shown.

### WT1 regulates microRNA levels

WT1 interaction with both the pre- as well as the mature microRNA suggest that the WT1-RNA complex could be involved in either microRNA targeting or microRNA processing. Since in an earlier study, we used FLASH to identify WT1 interaction with RNA hybrids (Bharathavikru et al., 2017), showing preferential enrichment towards the 3’ UTR of the transcript, we decided to assess if any significant WT1-associated miRNA-mRNA 3’ UTR hybrids were present in the FLASH dataset. If there were miRNA-mRNA hybrids (indicating a role in microRNA targeting), the candidate mRNAs should exhibit an upregulation of expression when comparing their transcriptome with the Wt1 knockdown cells (*Wt1 kd*) (Bharathavikru et al., 2017). Although microRNA-mRNA hybrids were identified, these did not show any change in expression; only a few were downregulated in expression (**Table S1**).

These observations suggest that WT1-microRNA interaction was most likely to be involved in the microRNA-processing pathway. Hence, we decided to investigate whether there were any changes in miRNA levels in WT1 expressing cells. Since multiple let-7 microRNAs were found to interact with WT1 and let-7c was present in both RIP and FLASH datasets (Bharathavikru et al., 2017), it was selected as one of the representative microRNAs for most of the validation studies. Northern blotting analysis of cells where WT1 is depleted (*Wt1* knockdown M15 cells or *Wt1* knockout embryonic stem (ES) cells) showed a downregulation of mature miRNA let-7c levels (**Panel I, Fig. 2A and 2B**) and an increase in the pre-microRNA levels (**Panel II, Fig. 2A and 2B)**, suggesting that, in the absence of WT1, the microRNA processing pathway is defective from the dicer node of the pathway. In general, we observed a decrease in mature microRNAs in cells lacking WT1 (**Fig.S2A**), which was validated using individual microRNA specific primers for a representative subset.

**Figure 2:**
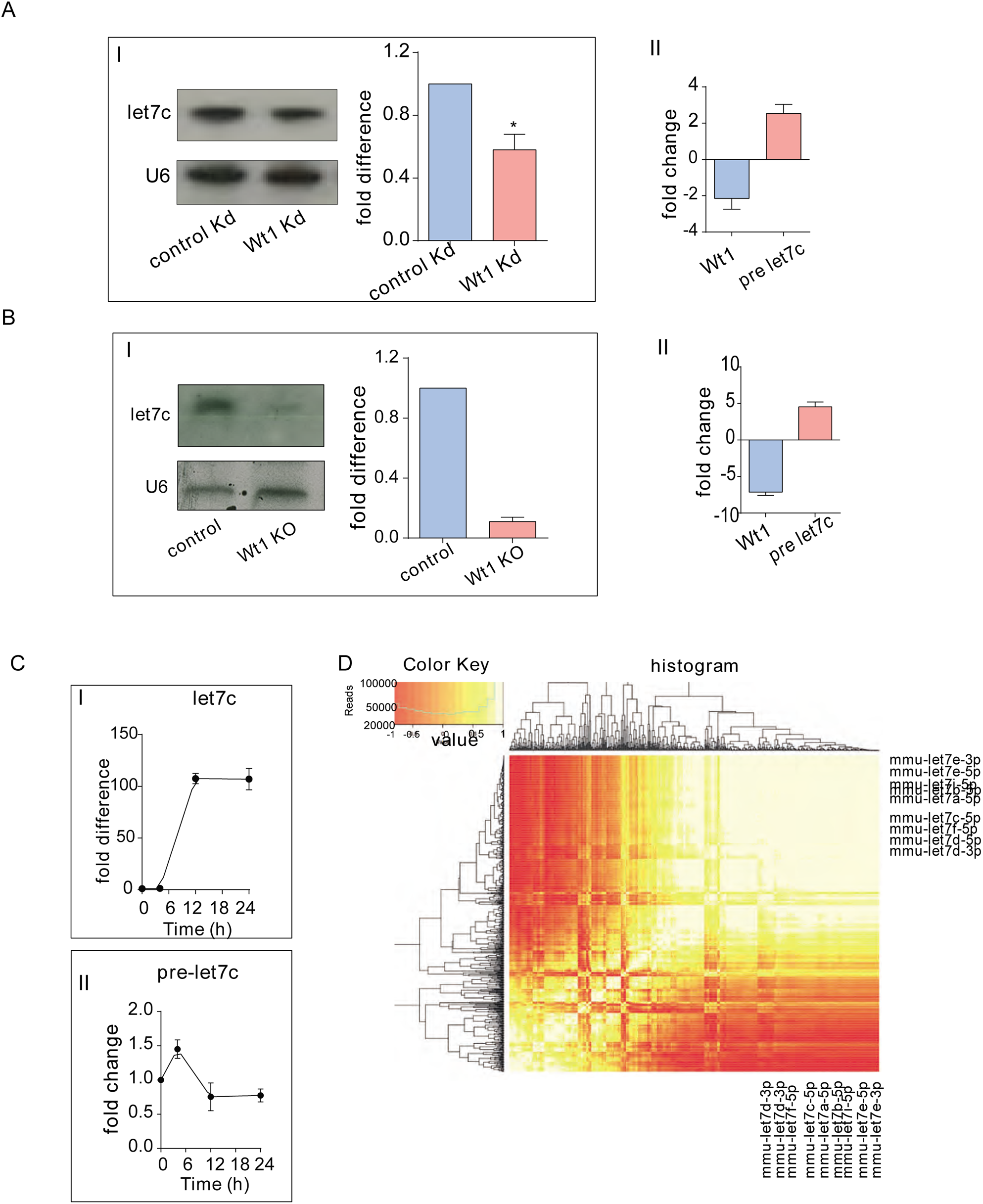
WT1 regulates microRNA levels. A) Northern blotting analysis and quantification (p<0.01) of mature let-7c expression (Panel I), compared to U6 RNA loading control and qPCR analysis of premiRNA let-7c (Panel II) in M15 cells compared to *Wt1* knockdown. B) Northern blotting analysis and quantification of mature let-7c expression (Panel I), compared to U6 loading control and qPCR analysis of premiRNA let-7c levels (Panel II) in ES cells compared to *Wt1* knockout cell line. C) Quantification of let-7c expression by northern blotting (Panel I), normalized to U6 RNA loading control, represented as the fold difference relative to time 0; qPCR analysis of pre-miRNA let-7c levels (Panel II) in RNA isolated from *Wt1* KO ES cells transfected with mcherry fused WT1, induced with doxycycline (1µg) for 0, 4, 12 and 24 hours. D) Heatmap of pairwise correlations of WT1 expression in four conditions (Ku, D12, D24 and ERA, see main text) to 637 expressed mature miRNA. A large proportion of miRNA have highly correlated expression patterns (clustered top right) and this population includes let-7s.

Since WT1 regulation of microRNA let-7 was affected more in the ES cells, we decided to investigate whether WT1 itself could regulate microRNA levels in these cells. Knockout ES cells were transiently transfected with a plasmid expressing the +exon5/+KTS isoform of WT1 (**Fig. S2B**). In the plasmid, the expression of WT1 is under the influence of doxycycline (dox) regulation, hence, dox treatment was used to titrate the levels of WT1. Induction of WT1 was confirmed by western blotting (**Fig.S2C**). Accordingly, northern blots revealed that, within 4 hours of dox induction, there was a slight increase in the let-7 microRNA levels that further increased upon 12 hours of dox treatment and saturated by 24 hours of treatment (**Fig. 2C, panel I**) with a corresponding downregulation of premature microRNA let-7c, detected by qPCR (**Fig. 2C, Panel II**). To visualise the overall pattern of microRNA expression changes, we performed microRNA sequencing and compared the expression of 637 detectable microRNAs across four samples with varying expression levels of WT1: knockout ES cells without dox (KU), dox 12 hours (D12), dox 24 hours (D24) and ES cells treated with retinoic acid for 5 days (ERA) (**Fig. 2D**). We correlated the expression of each microRNA with WT1 expression across the four samples and observed a highly correlated cluster of miRNAs that included let-7 miRNAs (**Fig. 2D**). Indeed, the expression of 9 detectable mature let-7 transcripts appeared to increase with higher WT1 levels (i.e. from Ku to ERA, **Fig.S2D**). It can be concluded that the transient induction of WT1 is sufficient to increase the microRNA levels, and especially the levels of microRNA let-7, in *Wt1* knockout ES cells.

Steady-state levels of let-7 microRNAs are regulated by the RNA binding protein LIN28 (Tsialikas and Romer-Seibert, 2015, Viswanathan et al., 2008). In ES cells, WT1 expression increases during a 5-day retinoic acid (RA) time course (Bharathavikru et al., 2017, Spraggon et al., 2007). During the same time period of differentiation, let-7 levels also increase with a concomitant decrease in LIN28 expression. Therefore, we decided to investigate whether LIN28 plays a role in WT1-mediated microRNA processing. As expected, *Wt1* levels increase across the 5 days of RA treatment in the wildtype cells, whereas the KO ES cells do not respond in a similar manner (**Fig.S2E**). *Lin28* levels, however, showed minimal differences between wildtype and knockout cells across the same time course (**Fig. S2E**).

### WT1 interacts with the microRNA pathway components

Earlier studies have shown accumulation of pre-microRNAs when MIRPG is deleted (Lee et al., 2004). Hence, western blotting analysis of the microRNA processing pathway components including DROSHA, DDX5 and AGO proteins was performed. We did not see any significant changes of these proteins, corresponding to the presence or absence of WT1 both in ES cells and M15 cells (**Fig. 3A**). Thus, the observed defect in the microRNA processing (accumulation of pre-processed miRNA), in the absence of WT1, is not due to any alterations in the microRNA processing machinery and is independent of the LIN28 axis.

**Figure 3:**
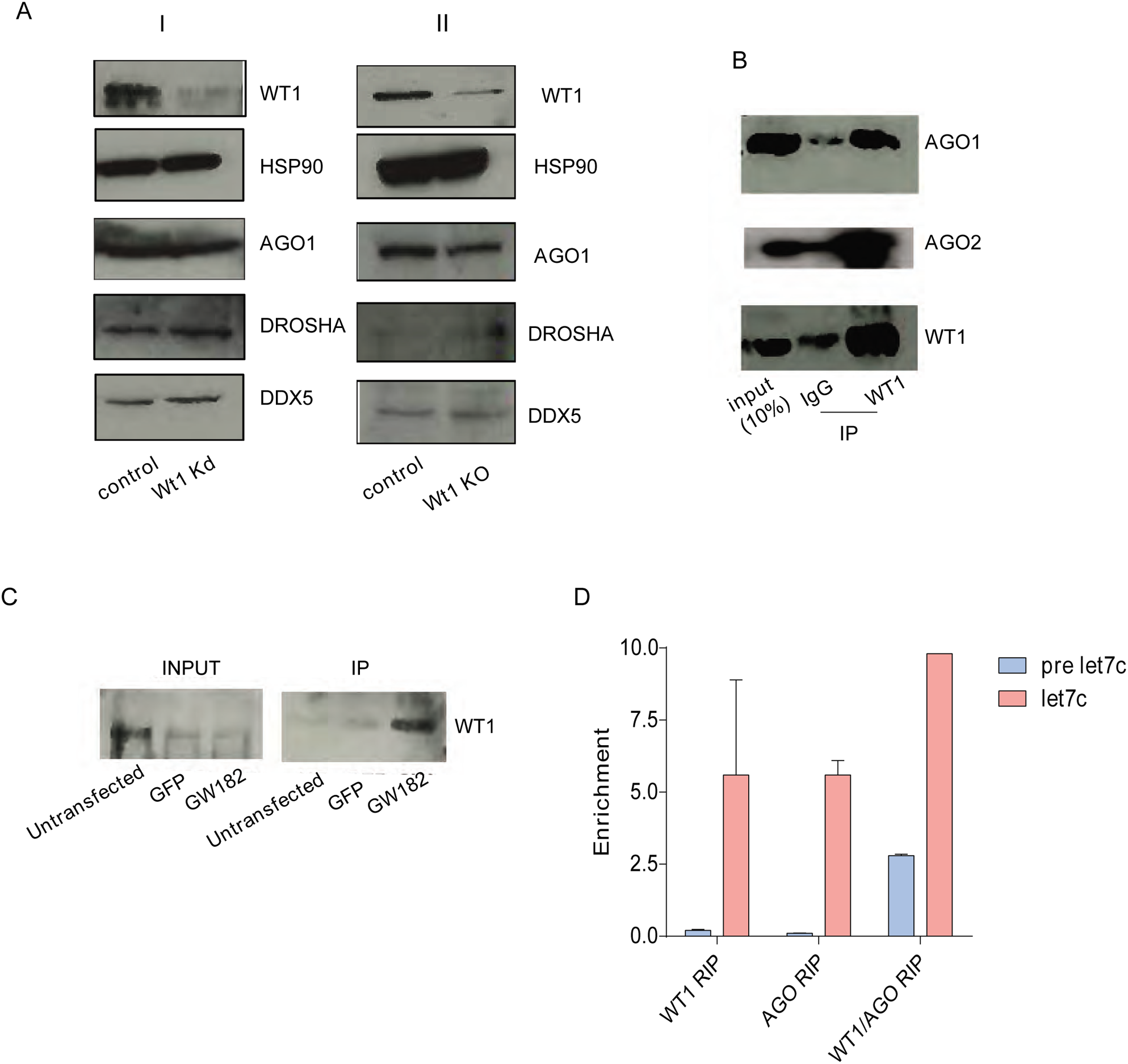
WT1 interacts with the microRNA pathway components. A) Western blotting analysis of M15 and ES cells compared to the respective knockdown and knockout cell lines, showing no observable changes in the levels of the microRNA processing pathway proteins (AGO1, DROSHA,DDX5). HSP90 is used as a loading control. WT1 levels depict the knockdown and knockout conditions respectively. B) Immunopulldowns using WT1 antibody and control IgG antibody, assessed for interacting proteins of the microRNA processing pathway (AGO1/2), compared to the respective input (10%) samples. C) GFP-GW transfected lysates (lane 3) along with untransfected (lane 1) and GFP alone (lane 2) transfected lysates were subjected to immunopulldowns against GFP antibody and immunoblotted against WT1 antibody, showing interaction of GW182 with WT1. D) qPCR analysis of RNA immunopulldowns to assess microRNA levels in WT1 bound RNA, AGO2 bound RNA and WT1-AGO bound RNA conditions. Fold enrichment was calculated comparing WT1 RIP to corresponding IgG RIP conditions.

In order to gain insights into whether WT1 regulates microRNA processing directly or through indirect mechanisms, Immunopulldowns (IPs) were used to assess if any of the microRNA biogenesis pathway proteins (from the DICER step onwards), are WT1 interacting proteins (**Fig. 3B**). It has been previously shown that WT1 interacts with the microRNA pathway components DICER and AGO2 (Akpa et al., 2016). WT1 interaction with AGO1 and AGO2 was confirmed in the mesonephric cells (**Fig. 3B**). The GW protein family (TNRC6a/b/c), which act as adaptor proteins for AGO interactions (Pfaff et al., 2013, Liu et al., 2018), was also assessed. GFP tagged GW182 was overexpressed in cells expressing WT1 and subjected to IP with antibody against GFP and immunoblotted against WT1 antibody which showed a positive interaction with WT1 (**Fig. 3C**). Thus, the microRNA processing pathway proteins AGO 1/2, GW adaptor protein were found to interact with WT1 implying an association of WT1 with the microRNA processing machinery.

The WT1-AGO interaction, suggests that the WT1-AGO complex will have an increased affinity to the mature microRNAs. In order to confirm this hypothesis, we performed double RIP experiments where cells were subjected to an initial round of RNA immunopulldown against WT1 and the eluted WT1-RNA bound complex was processed for a second RNA immunopulldown using an antibody against AGO2. WT1 interaction with pre-miRNA let-7 was considerably higher than the AGO- pre-miRNA association. Individual RIP of AGO and WT1 showed similar enrichment of mature let-7, whereas the mature let-7 bound to the WT1-AGO complex showed a much higher enrichment (**Fig. 3D**). The WT1-AGO bound RNA was subjected to a low depth FLASH analysis and the single reads were found to be significantly different from the WT1 bound RNA complex. The WT1-AGO interacting RNA component had a larger proportion of noncoding RNAs of which the microRNA component was more than 60% (**Fig. S3A**). The individual plots of let-7 microRNAs identified in the WT1-AGO bound RNA cluster in comparison to the WT1 bound RNA cluster showed increased reads with the WT1-AGO complex (**Fig. S3B**).

### The WT1-microRNA let-7-*Igf1r* axis in the kidney mesenchyme

The physiological relevance of the regulation of let-7 microRNA levels would reflect on the expression of the mRNA targets that are regulated by let-7. Let-7 microRNA levels are downregulated in the absence of WT1, indicating a possible upregulation of its corresponding target genes. To investigate this, we analysed the levels of a representative subset of TARGETSCAN predicted let-7 targets in *Wt1* knockdown M15 mesonephric cells. Two of the predicted let-7 targets, *Lin28* and, more strikingly, *Igfr1* showed significant upregulation upon *Wt1* knockdown (**Fig. 4A**), whereas the other predicted targets were either unaffected or downregulated. The likely explanation for this is that most of these are validated WT1 transcriptional targets (eg: *Co1a1, Igf2*), positively regulated by WT1 (Motamedi et al.,2014), as well as other downstream pathways regulated by WT1 transcriptional targets.

**Figure 4:**
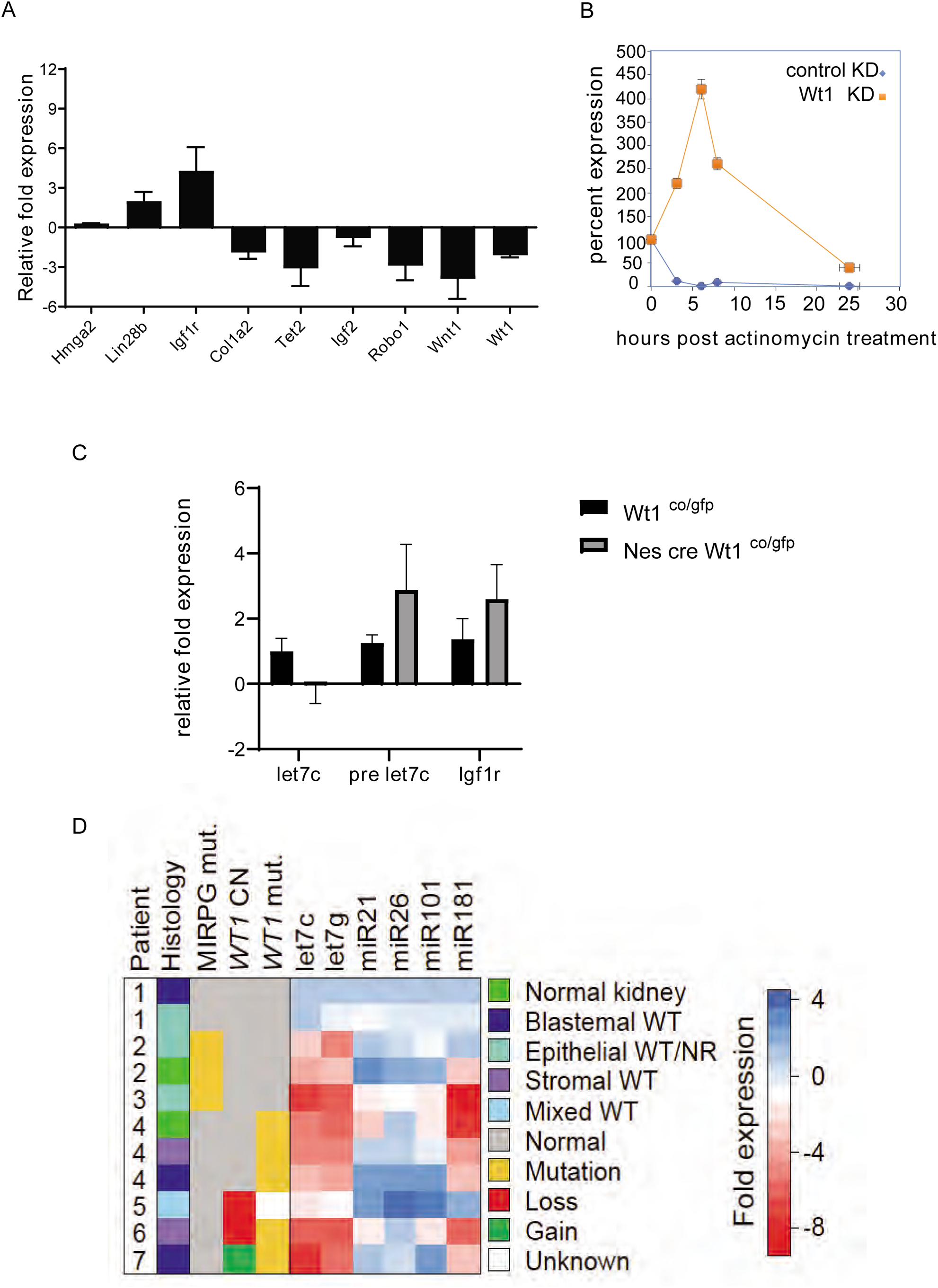
WT1 regulates microRNA let-7 levels in developing as well as diseased kidneys. A) qPCR analysis of TARGETSCAN-predicted let-7 targets, in the presence (*lacz kd*) or absence of WT1 (*Wt1 kd*), represented as fold change in expression, normalized to 18s RNA expression. B) RNA stability measurements, depicted as percent RNA of *Igf1r* mRNA following actinomycin treatment for the indicated time in control cells and cells deleted for *Wt1.* C) qPCR analysis of pre-miRNA let-7c, miRNA let-7c and *Igf1r* levels in E13.5, *Wt1* deficient kidney mesenchyme compared to litter matched *gfp* control. The expression is represented as relative fold expression compared to the control samples after normalizing to 18s RNA levels. D) For seven patients, the histology, mutational status for MIRPG, copy number for WT1 (MLPA) and mutational status for WT1 is displayed. MIRPG mutants included DGCR8 (patient 2) and DICER (patient 3). *WT1* mutations included 1568C>CT, 458R>R/X, 948_949het_insG (patient 4), Ins CGTAGAC (patient 6) and NM_000378:c.G208A:p.G70S (patient 7). For the microRNAs displayed, the fold change in expression (based on qPCR) is shown, relative to patient sample 1, with normal expression of *MIRPG* genes and *WT1*.

We have previously shown that WT1 stabilizes mRNAs (Bharathavikru et al., 2017). Many of the let-7 targets did not show any appreciable 3’ UTR interaction, nor change in expression with WT1. However, *Igf1r*, which does not interact with WT1, showed an upregulation in expression in the absence of WT1 (**Fig. 4A**). To confirm that the changes in *Igf1r* levels are indeed through mechanisms other than transcription, an RNA stability assay was performed in cells treated with actinomycin which were assessed for the RNA levels of *Igf1r* after different time points, which revealed that the upregulation was mostly post-transcriptional in nature (**Fig. 4B**).

WT1-mediated regulation of the microRNA let-7 pathway was observed in two different cell lines expressing WT1. To confirm whether this was relevant in a physiological context, *Wt1* was deleted in the kidney mesenchyme using the *Nestin-cre Wt1* conditional mouse line and the *Wt1 GFP* knockin model (Essafi et al., 2011). Embryonic day E13.5, metanephric mesenchyme cells from *gfp* control (GFP positive, *Wt1*^*co/gfp*^) and *Wt1* deficient (GFP positive, *Nes cre Wt1*^*co/gfp*^) genotypes were sorted by FACS to isolate the populations of GFP positive cells. RNA purified from the control and mutants was assessed for microRNA levels. In the mutants, mature let-7 microRNA expression was downregulated whereas, pre-mature miR let-7 was significantly upregulated consistent with the observation in cell lines (**Fig. 4C**). Correspondingly, in *Wt1*-deleted kidney mesenchyme, the *Igf1r* levels show significant upregulation as compared to the controls (**Fig. 4C**), suggesting that the WT1-let-7-*Igf1r* axis is functional in the tissue context.

*WT1* is frequently mutated in the paediatric kidney cancer, WT, and several studies have shown that microRNA-processing pathway genes are also mutated with an associated decrease in mature microRNA levels (Wegert et al., 2015, Gadd et al., 2017, Ludwig et al., 2016, Torrezan et al., 2014, Rakheja et al., 2014). To see whether *WT1* mutations may also impact microRNA expression levels, we performed a preliminary small-scale analysis with patient samples. The expression levels of six microRNAs were assessed by qPCR in a representative panel of WT and normal kidney samples with *WT1* mutations or an aberrant gene copy number, as well as samples with MIRPG mutations (*DICER* or *DGCR8*) as positive controls (**Fig. 4D**). In comparison to unmutated WTs, MIRPG samples displayed altered microRNA expression levels, including downregulation of let-7c, let-7g and miR181 (**Fig. 4D**). Supporting the role of WT1 in microRNA processing, samples with *WT1* mutation showed remarkably similar microRNA expression levels to MIRPG mutated samples (**Fig. 4D**). Due to the FFPE samples used for analysis, the expression of the pre microRNAs could not be assessed. All the observations suggest that WT1 mediated regulation of microRNA let-7 expression is relevant both in the context of mouse nephrogenesis as well as in WT. Thus, it can be concluded that WT1 interacts with and facilitates the processing of a subset of microRNAs, in particular the let-7 family, leading to the modulation of the IGF signalling pathway **(Fig.S4)** through IGF1R levels.

## Discussion

In this study, we show that WT1 interacts with and regulates microRNAs, especially the let-7 family. This interaction was of a bimodal pattern, suggesting a possible interaction of WT1 with the stem region of the stem-loop structure. WT1 has already been shown to interact with 3’ UTR of mRNAs especially with secondary structures, stabilizing them (Bharathavikru et al., 2017), a significant proportion of these mRNAs regulate developmental processes. WT1 interacts with microRNA let-7 and facilitates its processing, suggesting a negative modulatory role in RNA turnover. The transient transfection of *Wt1* in cells lacking its expression was sufficient to increase the microRNA levels. The results from the loss of function and gain of function studies reported here, support a posttranscriptional role for WT1 in microRNA processing. The global sequencing experiment revealed different categories of correlation with microRNA and WT1 expression, with the largest category of correlated clusters that included several members of the let-7 microRNA family. Within these correlation clusters, it is also possible that there are microRNAs transcriptionally regulated by WT1, and further investigations are required to identify them. WT1 as a transcription factor is known to activate and repress gene expression (Hastie, 2017). Further studies on WT1-RNA interaction could possibly highlight such a dual regulatory role in the RNA context as well.

WT1 interacts with the microRNAs as well as microRNA processing pathway protein components. The observation in **Fig.1D** points to a role for WT1 after the Dicer step in the microRNA pathway. The presence of the WT1-AGO complex, also suggests a role in the final steps of the microRNA pathway. Thus, WT1 role in the microRNA pathway seems to be in enabling the pre mature miRNA to be assembled into the functional RNA induced silencing complex. WT1 regulates gene expression through transcriptional and post transcriptional mechanisms. This is also evident when let7 targets are examined in the absence of *Wt1* (**Fig.4A**), when most of the genes show downregulation of expression. This can be attributed to the transcriptional role of WT1 and additional feedback loops that regulate microRNA pathways, which requires a detailed analysis.

The let-7 microRNA is regulated by the RBP, LIN28, which is essential for maintaining homeostasis. Lin 28 overexpression has been correlated to tumorigenesis (Viswanathan et al., 2009). In the cells that express WT1, we observe minimal changes in the LIN28 levels, however, there is a marked decrease in the let-7 levels. Since, the misexpression of *lin28* or let-7 impacts cellular differentiation, *WT1* mutations as well as the *MIRPG* mutations, all lead to a decrease in the let-7 levels, thus influencing nephron progenitor differentiation. In agreement with this, is the recent finding that the lin28-let-7 axis and the timing of let-7 expression regulates nephrogenesis (Yermalovich et al., 2019). The RNA degradation pathway component, *DIS3L2*, that is involved in the lin28-let-7 axis, has also been previously shown to be one of the candidate mutations that leads to WT. Thus, perturbation of the microRNA let-7-lin28 pathway seems to be one of the key pathways that result in WT. The lin28-let-7-IGF is a well-established regulatory node in glucose metabolism (Zhu et al., 2011). The cellular and metabolic events that could influence the role of WT1 in the microRNA processing pathway also need to be investigated. It is quite possible that such tissue specific factors would respond to different signals, as opposed to general transcription factors and ubiquitously expressed RNA binding proteins (Treiber et al., 2017).

Let-7 microRNA has been implicated in fine-tuning the process of differentiation. In line with this observation, is the study where LIN28 over expression under the *Wt1* promoter in early progenitor cells led to WT (Urbach et al., 2014). To counter this upregulated lin28 levels, the exogenous introduction of let-7g microRNA led to tumour abrogation. Thus, the let-7 microRNA pathway seems to be protective in the context of WT. On the other hand, *Dis3l2* and *Drosha* mutations have been tested in mouse models (Hunter et al., 2018, Kruber et al., 2019), suggesting that, although they are tumour promoting, they do not result in WT. This is also evident in our study where the early kidney mesenchyme already shows an upregulation of the pre-processed form of microRNAs. WT is composed of heterogeneous cell populations that are part of the nephrogenesis pathway, and it has been hypothesized that there are cancer stem cells that lead to the manifestation of WT (Shukrun et al., 2014). In this context, the microRNA pathway, especially the lin28-let-7 axis, could provide significant insight into the causative mechanisms and pathways that underlie WT. There are multiple events that lead to WT, including the perturbation of the IGF2 signalling pathway (Bharathavikru and Hastie, 2018). This occurs through genetic and/or epigenetic mechanisms including paternal disomy or loss of imprinting of the maternal allele. Recently, it has been shown that microRNA dysregulation can also relate to PLAG1 overexpression that leads to IGF2 upregulation, thus contributing to WT (Chen et al., 2018). Additionally, DIS3L2 has also been shown to be involved in the tumour biology by upregulating IGF2 expression (Hunter et al., 2018). In agreement with this augmented IGF expression, we observe changes in *Igf1r* levels as a downstream event of let-7 downregulation. WT1 regulates the RNA stability of IGFBPs, including IGFBP5. Both IGFBPs and IGF receptors bind to the growth hormone IGF: IGFBPs sequester IGF and the IGF-IGF receptor interaction potentiates the signalling pathway (Kim et al., 2009, Chao et al., 2008). In the absence of WT1, IGFBPs are downregulated (Bharathavikru et al., 2017) while IGF1R is upregulated, suggesting an increase in the IGF signalling events (**Fig. S4**). Previously, it has been shown that in WT, genetic and/or epigenetic mechanisms lead to increased IGF2 expression, and IGF1R is also upregulated in WT through copy number gain and linked to the possibility of relapse (Natrajan et al., 2006). We now show that, in addition to these events, WT1-mediated let-7 microRNA regulation and the downstream IGF1R upregulation also seems to be another perturbed node that, enhances the IGF signalling pathway and contributes towards the manifestation and pathology of WT. This needs to be addressed in a separate study, with an increased number of patient samples across different mutations and compared for the perturbation of the WT1-let-7-IGF1R axis. The microRNA pathway has been recently explored for early diagnostic purposes as well as therapeutic targeting. Circulating microRNAs can be detected from body fluids at very initial stages of diseases and thus help with early detection (Wang et al., 2018). However, the most promising therapeutic strategy seems to be the IGF signalling pathway. Incidentally, IGF1R has been tested as a therapeutic target in several cancers (Crudden et al., 2015). Our study not only provides a mechanistic insight into the causes of WT but also suggests potential diagnostic and therapeutic strategies involving the microRNA let-7 and the IGF signalling pathway.

## Methods

### FLASH and RIPseq

were performed as described earlier in Bharathavikru et al., 2017. Briefly, cells were crosslinked either by UV followed by formaldehyde or formaldehyde alone and subjected to Immunopulldown of WT1-RNA complexes through an antibody against WT1. The immune complexes thus precipitated where subjected to adaptor ligation and sequencing to identify the RNA subtypes that are associated with WT1.

### miRNA purification

Total RNA from cell lines was isolated as per the manufacturers’ protocols using the Qiagen mRNeasy kit. Total RNA from kidney mesenchyme was purified using TRIZOL and ethanol precipitation. RNA was quantified using nanodrop and agilent bioanalyzer to verify the concentration as well as the purity of isolated RNA. WT patient FFPE samples were processed for xylene deparaffinization, followed by miRNA purification following the manufacturers’ protocol in the Qiagen FFPE miRNeasy kit.

### miRNA northern blotting protocol

miRNA was either selectively purified or total mRNA was used with probes specific for miRNAs to facilitate detection. DNA oligo was designed by using the template for antisense probe, synthesized oligo was resuspended in nuclease free water so as to obtain a 100µM stock solution. This oligo template was hybridized to T7 promoter primer and heated to 70°C for 5mins followed by room temperature incubation for 5 minutes to facilitate hybridization. Hybridized oligo was processed for klenow filling using exo-klenow, incubated at 37°C for 30mins followed by the transcription reaction catalysed by T7 RNA polymerase in the presence of labelled UTP.

For the process of northern blotting, 10% Urea-PAGE was used for electrophoresing 10µg of total RNA samples, were blotted onto Nylon membrane for 1 hour using TBE as the buffer. After transfer, the membrane was UV crosslinked and processed for prehybridization for atleast 4 hours at 40°C in 10 ml of hybridization buffer. Following prehybridization, the labelled probe was added. After an overnight hybridization at 40°C, the samples were washed with 50ml of pre-warmed wash buffer followed by exposure to phosphorimager and an autoradiogram. Quantification of signals was performed using Image J analysis software, wherein U6 RNA levels were used as the loading control to normalize and calculate the expression difference across the test samples.

### Small RNA sequencing

Total RNA equivalent to 1µg was ligated with the 3’ RNA adapter with the help of the T4 RNA ligase 2 (deletion mutant) followed by ligation of the 5’ RNA adapter using T4 RNA ligase. The adapter ligated RNA was then subjected to reverse transcription with Superscript II RT followed by PCR amplification. The amplified cDNA was electrophoresed on a 6% PAGE gel. Mature microRNA, which is approximately 22 bp, gives an adapter ligated fragment of approximately 147 bp. A second band of 157 bp corresponding to piRNAs, other small noncoding RNA arising from 30 bp small RNAs was also excised together to give rise to the pool of small RNA. Additionally, the pre-processed forms of microRNA that result in longer fragments of approximately 160 bp to 300 bp were excised together for the pool of pre-processed small RNAs. The gel purified cDNA libraries were validated and sequenced on an Illumina platform.

### Dox inducible expression

Mcherry tagged *Wt1* construct was cloned into a plasmid with doxycycline inducible CAG promoter and sequence confirmed. Following transient transfection of *Wt1* KO embryonic stem cells with the above construct, dox induction was performed at different time points using 1µg/ml doxycycline. Cells were checked under the microscope at different time points to confirm mcherry expression. Induction of WT1 expression was confirmed by qRT-PCR analysis and western blotting. Based on the above results it was confirmed that dox inducible Wt1 expression was evident from 4 hours by qRT PCR wheareas the mcherry expression could be detected clearly under the microscope following an overnight induction of approximately 8 hours with 1µg/ml dox. *Wt1* KO ES cells were transiently transfected with the construct along with an mcherry alone expressing empty vector control. Following overnight transfection, the cells were induced with 1µg/ml dox and collected at the following time intervals, 0 hours, 4 hours, 12 hours and 24 hours of dox induction. Cells were always checked under a microscope for mcherry expression, the cells thus collected were processed for RNA isolation. Isolated RNA was quantified and subjected to cDNA synthesis for premicroRNA assessment as well as processed for northern blotting analysis to assess mature microRNA levels. The 12 hour and 24 hour post induction samples were also processed for small RNA sequencing. The sequenced miRNA was analysed as mentioned in Eminaga et al., 2013.

### qPCR for detection of miRNA

Premature microRNA was quantified by designing primers against the full length of the premature microRNA. cDNA synthesis was carried out using total RNA and random primers. This was then processed for qPCR analysis using SYBR green based detection methods. Fold change was calculated by normalizing to U6 snRNA or 18S rRNA and then comparing the control (atleast n=2) to the mutant lines (atleast n=3).

For, mature microRNA expression, the clontech mir-X miRNA first strand synthesis and SYBR kit was used which involves an initial polyadenylation reaction and synthesis. MicroRNA detection was carried out by amplifying the cDNA with a 3’ primer that recognizes the polyadenylated fragments and the specificity is brought about by a 5’ microRNA specific primer (that corresponds to the entire mature microRNA sequence) and SYBR green based detection. Calculations were carried out as explained above. This methodology was used for detection of mature microRNA in the FACS sorted kidney mesenchyme tissue as well as the miRNA from the FFPE tissue.

### Subcellular RNA fractionation

The RNA fractionation was performed as per the manufacturers protocol using the Active motif, subcellular RNA isolation kit. Primer pairs specific for differentially localized RNAs were used to confirm the efficiency of fractionation. Data was normalized by comparing to the total RNA content from the same cells, and calculating the enrichment as percent input as follows, 100x(2 exp Ct total RNA – Ct RNA fraction). The above RNA samples were then subjected to immunopulldowns as described below.

### RNA Immunopulldown (RIP)

RIP was performed as described (Bharathavikru et al., 2017). Briefly, cells were formaldehyde crosslinked, and sonicated, followed by immuno pulldown with WT1 antibody. After overnight incubation and washes, samples were reverse cross linked, proteinase K treated and RNA was precipitated. Primer pairs specific for pre microRNAs and mature microRNAs were used with the clontech mir-X miRNA first strand synthesis and SYBR kit for detection of immune precipitated miRNAs.

### Double RNA IP

Cells were processed for RNA Immuno pulldowns by formaldehyde crosslinking, followed by lysis and sonication. Sonicated lysates were subjected to overnight IP with WT1 and IgG antibodies in RIP buffer. Bound immune complexes were washed with RIP lysis buffer and eluted with the same buffer, Eluted samples were processed for a second RNA immuno pulldown using AGO2 and IgG antibodies for 3 hours. After washes, RNA was precipitated after reverse crosslinking, and proteinase K digestion. Eluted RNA was quantified, reverse transcribed and processed for quantitative PCR analysis. Single pulldowns of Ago and WT1 RNA IP were compared to IgG single pulldown and quantified. Double immunopulldown of WT1-AGO was compared to the IgG double RNA pulldown respectively.

### Protein immunopulldowns (IPs)

Protein IP was performed as described in Bharathavikru et al., 2016. Briefly, cells were lysed in RIPA buffer and IPs with WT1 and control IgG antibodies was performed in the presence of RNase. Resulting immune complexes were then resolved on 10% SDS PAGE gels and processed for immunoblotting.

### Immunoblotting analysis

Protein lysates and immune complexes from IPs were electrophoresed on 10% SDS PAGE gels and probed with antibodies against WT1 (ab89901), AGO1 (CST 9388), AGO2 (CST C34C6), DICER (CST D38E7), GFP (ab6556), LIN28B (CST S7056), DROSHA (CST D28B1), Conformation specific secondary antibody (NEB L27AS).

### microRNA let-7 inhibitor treatment assay

miRvana let7 inhibitor and the negative control inhibitor were procured and resuspended to give a 10µM stock solution. Cells (M15 control knockdown and Wt1 knockdown mesonephric cell lines), were reverse transfected with the miRvana inhibitors at 30nM concentration and after an overnight incubation, were subjected to media change. These cells were collected approximately 36 hours post transfection, and processed for RNA isolation. RNA was reverse transcribed and further subjected to qPCR analysis as described above. The calculations of the fold change was done by comparing the negative inhibitor treated cells to the miRvana let7 inhibitor treatment and represented as 2^^-deldelCt^

### Statistical Analysis

All experiments were performed with atleast 3 biological replicates and technical replicates of each. Standard deviation was calculated to compute the error bars. Unpaired t-test was done to obtain the two-tailed p values, in graphpad prism.

## Acknowledgements

We thank Medical Research Council (Core Fund to Human Genetics Unit) and MRC grant MR/NR20405/1 for funding. The authors are grateful to Dr. Lee Spraggon, MSKCC (GFP GW182 plasmid), Dr. Sara Macias Ribela, UoE (microRNA experiment advice), Dr. Vinay Bulusu, (assistance with manuscript preparation), Dr.Cathy Duff (maintenance of transgenic mouse lines), Elizabeth Freyer (FACS facility).

## Author contributions

R.B, N.D.H, designed the study, R.B, J.C, J.S, S.A, analyzed data, R.B, J.S, J.C, G.P, V.S, performed experiments, R.B, J.C, KPJ, A.E, N.D.H, provided ideas and suggestions, R.B, K.P.J, N.D.H, supervised experiments, N.D.H, obtained funding, R.B, J.C, N.D.H, wrote the manuscript with inputs from all authors.

## References

1. Akpa, M. M. Iglesias, D. Chu, L. Thiébaut, A. Jentoft, I. Hammond, L. Torban, E. Goodyer, P.R. Wilms Tumor Suppressor, WT1, Cooperates with MicroRNA-26a and MicroRNA-101 to Suppress Translation of the Polycomb Protein, EZH2, in Mesenchymal Stem Cells. J Biol Chem. 291, 3785–3795 (2016).

2. Astuti, D. Morris, M.R. Cooper, W.N. Staals, R.H. Wake, N.C. Fews, G.A. Gill, H. Gentle, D. Shuib, S. Ricketts, C.J. Cole, T. van Essen, A.J. van Lingen, R.A. Neri, G. Opitz, J.M. Rump, P. Stolte-Dijkstra, I. Müller, F. Pruijn, G.J. Latif, F. Maher, E.R. Germline mutations in DIS3L2 cause the Perlman syndrome of overgrowth and Wilms tumor susceptibility. Nat Genet. 44, 277–284 (2012).

3. Bardeesy, N. Falkoff, D. Petruzzi, M.J. Nowak, N. Zabel, B. Adam, M. Aguiar, M.C. Grundy, P. Shows, T. Pelletier, J. Anaplastic Wilms’ tumour, a subtype displaying poor prognosis, harbours p53 gene mutations. Nat Genet. 7, 91–97 (1994)

4. Bharathavikru, R. Dudnakova, T. Aitken, S. Slight, J. Artibani, M. Hohenstein, P. Tollervey, D. Hastie, N. Transcription factor Wilms’ tumor 1 regulates developmental RNAs through 3’ UTR interaction. Genes Dev. 31, 347–352 (2017).

5. Bharathavikru, R. & Hastie, N.D. Overgrowth syndromes and pediatric cancers: how many roads lead to IGF2? Genes Dev. 32, 993–995 (2018).

6. Bharathavikru, R. & von Kriegsheim, A. WT1-Associated Protein-Protein Interaction Networks. Methods Mol Biol. 1467, 189–196. (2016).

7. Chang, H.M. Triboulet, R. Thornton, J.E. Gregory, R.I. A role for the Perlman syndrome exonuclease Dis3l2 in the Lin28-let-7 pathway. Nature 497, 244–248 (2013).

8. Chao, W. & D’Amore, P.A. IGF2: epigenetic regulation and role in development and disease. Cytokine Growth Factor Rev. 19, 111–120 (2008).

9. Charlton, J. & Pritchard-Jones, K. WT1 Mutation in Childhood Cancer. Methods Mol Biol. 1467, 1–14 (2016).

10. Chen, K.S. Stroup, E.K. Budhipramono, A. Rakheja, D. Nichols-Vinueza, D. Xu, L. Stuart, S.H. Shukla, A.A. Fraire, C. Mendell, J.T. Amatruda, J.F. Mutations in microRNA processing genes in Wilms tumors derepress the IGF2 regulator PLAG1. Genes Dev. 32, 996–1007 (2018).

11. Crudden, C. Girnita, A. & Girnita, L. Targeting the IGF-1R: The Tale of the Tortoise and the Hare. Front Endocrinol (Lausanne). 6, 64. Review. (2015).

12. Eminaga, S. Christodoulou, D.C. Vigneault, F. Church, G.M. & Seidman, J.G. Quantification of microRNA expression with next-generation sequencing. Curr Prot Mol Biol 4.17.1-4.17.4 (2013)

13. Essafi, A. Webb, A. Berry, R.L. Slight, J. Burn, S.F. Spraggon, L. Velecela, V. MartinezEstrada, O.M. Wiltshire, J.H. Roberts, S.G.E. Brownstein, D. Davies, J.A. Hastie, N.D. Hohenstein, P. A Wt1-Controlled chromatin switching mechanism underpins tissue-specific wnt4 activation and repression. Dev. Cell 21, 559–574 (2011).

14. Gadd, S. Huff, V. Walz, A.L. Ooms, A.H.A.G. Armstrong, A.E. Gerhard, D.S. Smith, M.A. Auvil, J.M.G. Meerzaman, D. Chen, Q.R. Hsu, C.H. Yan, C. Nguyen, C. Hu, Y. Hermida, L.C. Davidsen, T. Gesuwan, P. Ma, Y. Zong, Z. Mungall, A.J. Moore, R.A. Marra, M.A. Dome, J.S. Mullighan, C.G. Ma, J. Wheeler, D.A. Hampton, O.A. Ross, N. Gastier-Foster, J.M. Arold, S.T. Perlman, E.J. A Children’s Oncology Group and TARGET initiative exploring the genetic landscape of Wilms tumor. Nat Genet. 49, 1487–1494 (2017).

15. Hastie, N. D. Wilms’ tumour 1 (WT1) in development, homeostasis and disease. Development. 144, 2862–2872 (2017). Review.

16. Hu, Q. Gao, F. Tian, W. Ruteshouser, E.C. Wang, Y. Lazar, A. Stewart, J. Strong, L.C. Behringer, R.R. Huff, V. Wt1 ablation and Igf2 upregulation in mice result in Wilms tumors with elevated ERK1/2 phosphorylation. J Clin Invest. 121,174–183 (2011)

17. Hunter, R.W. Liu, Y. Manjunath, H. Acharya, A. Jones, B.T. Zhang, H. Chen, B. Ramalingam, H. Hammer, R.E. Xie, Y. Richardson, J.A. Rakheja, D. Carroll, T.J. Mendell JT. Loss of Dis3l2 partially phenocopies Perlman syndrome in mice and results in up-regulation of Igf2 in nephron progenitor cells. Genes Dev. 32, 903–908 (2018).

18. Kim, S.Y. Toretsky, J.A. Scher, D. Helman, L.J. The role of IGF-1R in pediatric malignancies. Oncologist. 14, 83–91 (2009).

19. Kruber, P. Angay, O. Winkler, A. Bösl, M.R. Kneitz, S. Heinze, K.G. Gessler, M. Loss or oncogenic mutation of DROSHA impairs kidney development and function, but is not sufficient for Wilms tumor formation. Int J Cancer. 144, 1391–1400 (2019).

20. Lee, Y.S. Nakahara, K. Pham, J.W. Kim, K. He, Z. Sontheimer, E.J. Carthew, R.W. Distinct roles for Drosophila Dicer-1 and Dicer-2 in the siRNA/miRNA silencing pathways. Cell. 117, 69–81 (2004).

21. Liu, J. Liu, Z. & Corey, D.R. The Requirement for GW182 Scaffolding Protein Depends on Whether Argonaute Is Mediating Translation, Transcription, or Splicing. Biochemistry. 57, 5247–5256 (2018)

22. Ludwig, N. Werner, T.V. Backes, C. Trampert, P. Gessler, M. Keller, A. Lenhof, H.P. Graf, N. Meese, E. Combining miRNA and mRNA Expression Profiles in Wilms Tumor Subtypes. Int J Mol Sci. 17, 475-.(2016).

23. Motamedi, F.J. Badro, D.A. Clarkson, M. Lecca, M.R. Bradford, S.T. Buske, F.A. Saar, K. Hübner, N. Brändli, A.W. Schedl, A. WT1 controls antagonistic FGF and BMP-pSMAD pathways in early renal progenitors. Nat Commun. 5, 4444-(2014).

24. Natrajan, R. Reis-Filho, J.S. Little, S.E. Messahel, B. Brundler, M.A. Dome, J.S. Grundy, P.E. Vujanic, G.M. Pritchard-Jones, K. Jones, C. Blastemal expression of type I insulin-like growth factor receptor in Wilms’ tumors is driven by increased copy number and correlates with relapse. Cancer Res. 66, 11148–11155 (2006).

25. Niksic, M. Slight, J. Sanford, J.R. Caceres, J.F. Hastie, N.D. The Wilms’ tumour protein (WT1) shuttles between nucleus and cytoplasm and is present in functional polysomes. Hum Mol Genet. 13:463–471 (2004).

26. Palculict, T.B. Ruteshouser, E.C. Fan, Y. Wang, W. Strong, L. Huff, V. Identification of germline DICER1 mutations and loss of heterozygosity in familial Wilms tumour. J Med Genet. 53, 385–388 (2016).

27. Pfaff, J. & Meister, G. Argonaute and GW182 proteins: an effective alliance in gene silencing. Biochem Soc Trans. 41, 855-860. Review. (2013).

28. Rakheja, D. Chen, K.S. Liu, Y. Shukla, A.A. Schmid, V. Chang, T.C. Khokhar, S. Wickiser, J.E. Karandikar, N.J. Malter, J.S. Mendell, J.T. Amatruda, J.F. Somatic mutations in DROSHA and DICER1 impair microRNA biogenesis through distinct mechanisms in Wilms tumours. Nat Commun. 5, 4802–4810 (2014).

29. Shukrun, R. Pode-Shakked, N. Pleniceanu, O. Omer, D. Vax, E. Peer, E. Pri-Chen, S. Jacob, J. Hu, Q. Harari-Steinberg, O. Huff, V. Dekel, B. Wilms’ tumor blastemal stem cells dedifferentiate to propagate the tumor bulk. Stem Cell Reports. 3, 24–33 (2014).

30. Spraggon, L. Dudnakova, T. Slight, J. Lustig-Yariv, O. Cotterell, J. Hastie, N. Miles, C. hnRNP-U directly interacts with WT1 and modulates WT1 transcriptional activation. Oncogene. 26, 1484–1491 (2007).

31. Torrezan, G. T. Ferreira, E.N. Nakahata, A.M. Barros, B.D. Castro, M.T. Correa, B.R. Krepischi, A.C. Olivieri, E.H. Cunha, I.W. Tabori, U. Grundy, P.E. Costa, C.M. de Camargo, B. Galante, P.A. Carraro, D.M. Recurrent somatic mutation in DROSHA induces microRNA profile changes in Wilms tumour. Nat. Commun. 5, 4039–4049 (2014).

32. Treiber, T. Treiber, N. Plessmann, U. Harlander, S. Daiß, J.L. Eichner, N. Lehmann, G. Schall, K. Urlaub, H. Meister, G. A Compendium of RNA-Binding Proteins that Regulate MicroRNA Biogenesis. Mol Cell. 66, 270-284.e13 (2017)

33. Tsialikas, J. & Romer-Seibert, J. LIN28: roles and regulation in development and beyond. Development. 142, 2397–2404 (2015).

34. Urbach, A. Yermalovich, A. Zhang, J. Spina, C.S. Zhu, H. Perez-Atayde, A.R. Shukrun, R. Charlton, J. Sebire, N. Mifsud, W. Dekel, B. Pritchard-Jones, K. Daley, G.Q. Lin28 sustains early renal progenitors and induces Wilms tumor. Genes Dev. 28, 971–982 (2014).

35. Viswanathan, S.R. Daley, G.Q. Gregory, R.I. Selective blockade of microRNA processing by Lin28. Science 320, 97–100 (2008).

36. Viswanathan, S. R. Powers, J.T. Einhorn, W. Hoshida, Y. Ng, T.L. Toffanin, S. O’Sullivan, M. Lu, J. Phillips, L.A. Lockhart, V.L. Shah, S.P. Tanwar, P.S. Mermel, C.H. Beroukhim, R. Azam, M. Teixeira, J. Meyerson, M. Hughes, T.P. Llovet, J.M. Radich, J. Mullighan, C.G. Golub, T.R. Sorensen, P.H. Daley, G.Q. Lin28 promotes transformation and is associated with advanced human malignancies. Nat Genet. 41, 843–848 (2009).

37. Wang, H. Peng, R. Wang, J. Qin, Z. & Xue, L. Circulating microRNAs as potential cancer biomarkers: the advantage and disadvantage. Clin Epigenetics. 23, 59. Review. (2018).

38. Wegert, J. Ishaque, N. Vardapour, R. Geörg, C. Gu, Z. Bieg, M, Ziegler, B. Bausenwein, S. Nourkami, N. Ludwig, N. Keller, A. Grimm, C. Kneitz, S. Williams, R.D. Chagtai, T. Pritchard-Jones, K. van Sluis, P. Volckmann, R. Koster, J. Versteeg, R. Acha, T. O’Sullivan, M.J. Bode, P.K. Niggli, F. Tytgat, G.A. van Tinteren, H. van den Heuvel-Eibrink, M.M. Meese, E. Vokuhl, C. Leuschner, I. Graf, N. Eils, R. Pfister, S. M. Kool, M. Gessler, M. Mutations in the SIX1/2 pathway and the DROSHA/DGCR8 miRNA microprocessor complex underlie high-risk blastemal type Wilms tumors. Cancer Cell. 27, 298–311 (2015).

39. Winter, J. Jung, S. Keller, S. Gregory, R.I. Diederichs, S. Many Roads to Maturity: MicroRNA Biogenesis Pathways and Their Regulation. Nat Cell Biol. 11, 228–234 (2009).

40. Yermalovich, A.V. Osborne, J.K. Sousa, P. Han, A. Kinney, M.A. Chen, M.J. Robinton, D.A. Montie, H. Pearson, D.S. Wilson, S.B. Combes, A.N. Little, M.H. Daley, G.Q. Lin28 and let-7 regulate the timing of cessation of murine nephrogenesis. Nat Commun. 10, 168–178 (2019).

41. Zhu, H. Shyh-Chang, N. Segrè, A.V. Shinoda, G. Shah, S.P. Einhorn, W.S. Takeuchi, A. Engreitz, J.M. Hagan, J.P. Kharas, M.G. Urbach, A. Thornton, J.E. Triboulet, R. Gregory, R.I. DIAGRAM Consortium; MAGIC Investigators, Altshuler, D. Daley, G.Q. The Lin28/let-7 axis regulates glucose metabolism. Cell 147, 81–94 (2011).

