## Supplementary Information for "Tumour suppressor WT1 regulates the let-7-*Igf1r* axis in kidney mesenchyme"

[1]MRC Human Genetics Unit, Institute of Genetics and Molecular Medicine, University of Edinburgh, Western General Hospital, Crewe Road (S), Edinburgh, EH4 2XU, [2]School of Cellular and Molecular Medicine, Faculty of Biomedical Sciences, University of Bristol, University Walk BS8 1TD, [3]Department of Oncology, Great Ormond Street Hospital for Children, NHS Foundation Trust, Great Ormond Street, London WC1N 3JH, UK.

[5]Present address: Department of Biological sciences, Indian Institute of Science Education and Research, Berhampur, Ganjam, Odisha 760010

**Figure S1: WT1 interacts with microRNAs**

(A) Schematic of WT1 interacting RNA identification and the subsequent validation approaches used in this study. (B) Graphical representation of noncoding and small RNA subtypes identified by RNA Immuno Precipitation (RIP) in mesonephric, M15 cells. (C) Northern blotting analysis of WT1 interacting let7c and U6 snRNA in input, IgG and WT1 RIP samples. M15 total RNA was used as a positive control. (D) Validation of subcellular compartmentalization using primer pairs specific for nuclear (blue) and cytoplasmic (orange) compartment.

**Figure S2: WT1 expression restores mature microRNA levels**

(A) qPCR analysis of a representative subset of microRNAs assessed for the pre-miRNA and mature microRNA levels (miR92, miR99 and miR125b) in Wt1 deleted M15 cells compared to control knockdown. (B) Construct used for WT1 expression in the knockout cells. (C) Western blotting analysis for the time course induction of WT1 expression upon dox addition compared to HSP90 loading control. (D) Boxplot representation of let7 levels for 9 mature let7 miRNA (let7a,b,c,d,e,f,i 5' transcripts and let7d,e 3' transcripts) in samples across wildtype ES cells and WT1 transfected KO ES cells, uninduced, induced with doxycycline for 12 hours and 24 hours. (E) Western blotting analysis of WT1, LIN28 across the 5 day RA time course, in Wt1 KO ES cells and ES cells. Histone H3 used as a loading control.

**Figure S3: WT1-AGO complex interacts with microRNAs.** (A) WT1-AGO bound RNA biotypes represented as pie charts. The bottom panel represents an enlarged version of the noncoding RNA biotypes. (B) The comparison of let7b/c microRNA profiles identified in the WT1-AGO microRNA bound subset compared to the WT1 bound microRNA subset.

**Figure S4: Schematic to explain role of WT1 in the let7-IGF1R axis in the kidney mesenchyme.** WT1 regulates Igfbp5 mRNA through regulating RNA stability. WT1 also regulates let7 microRNA processing and thus influences *Igf1r* levels. In the absence of WT1, the *Igfbp5* mRNA is less stable and *Igf1r* levels are increased, thus resulting in increased IGF signaling with potential implications in Wilms' Tumour manifestation.

**Table S1:** miRNA-mRNA hybrids identified from FLASH analysis and overlapping mRNA targets identified in the transcriptome analysis with their fold change upon *Wt1* knockdown.

**Table S2:** Oligo details

FIGURE S1

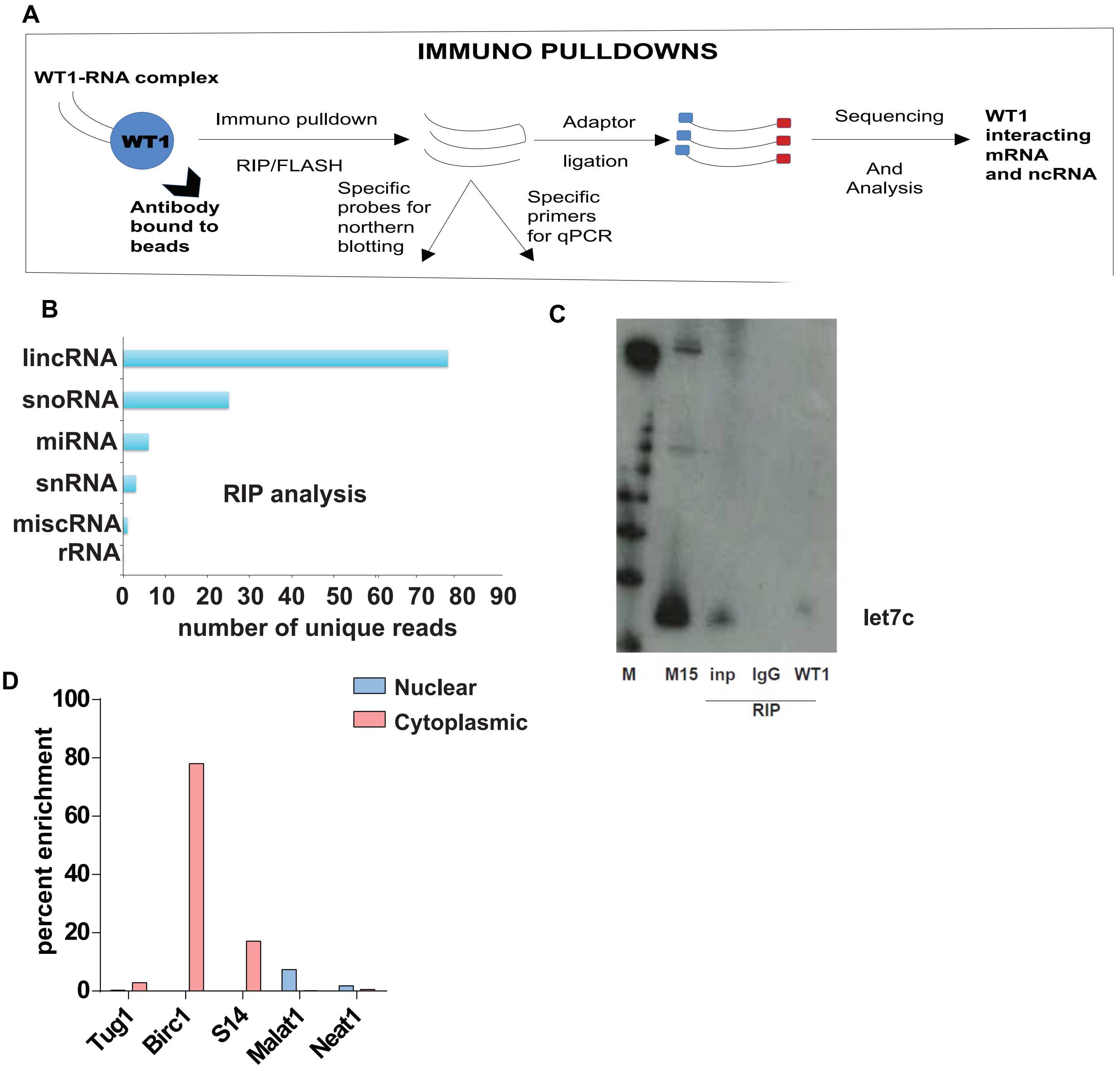


let7c

FIGURE S2

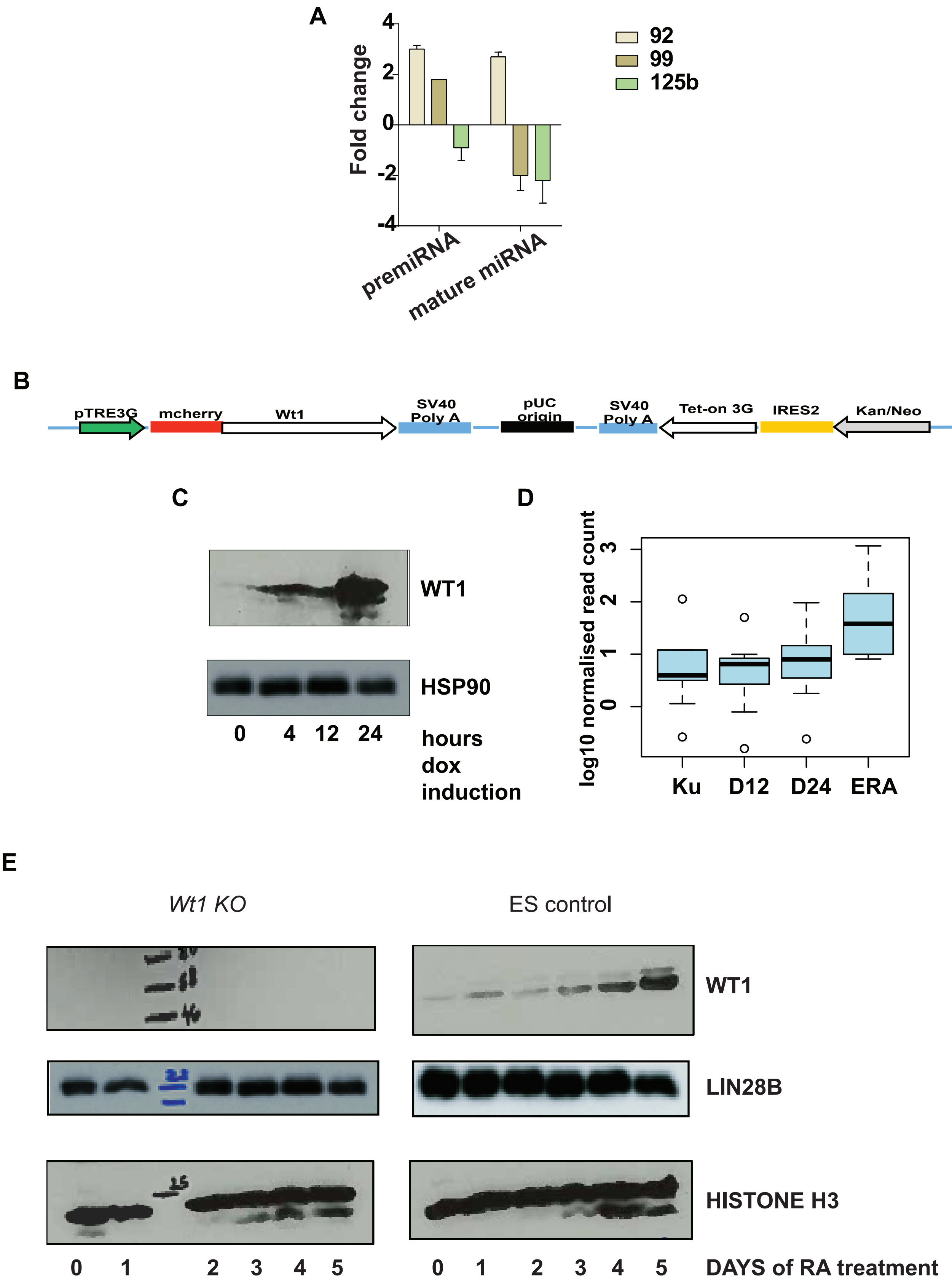
A
Fold change
4
2
0
-2
-4
premiRNA
mature miRNA
92
99
125b
B
pTRE3G
mcherry
Wt1
SV40 Poly A
pUC origin
SV40 Poly A
Tet-on 3G
IRES2
Kan/Neo
pSV40
C
WT1
HSP90
0
4
12
24
hours dox induction
D
log10 normalised read count
0
1
2
3
Ku
D12
D24
ERA
E
Wt1 KO
ES control
WT1
LIN28B
HISTONE H3
0
1
2
3
4
5
0
1
2
3
4
5
DAYS of RA treatment

A

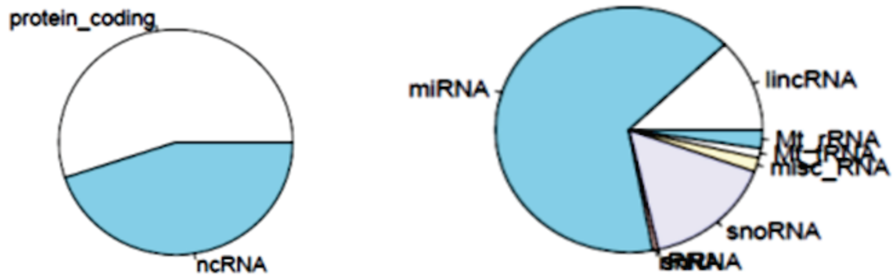
protein_coding
ncRNA
miRNA
lincRNA
Mt_rRNA
Mt_tRNA
misc_RNA
snoRNA

B

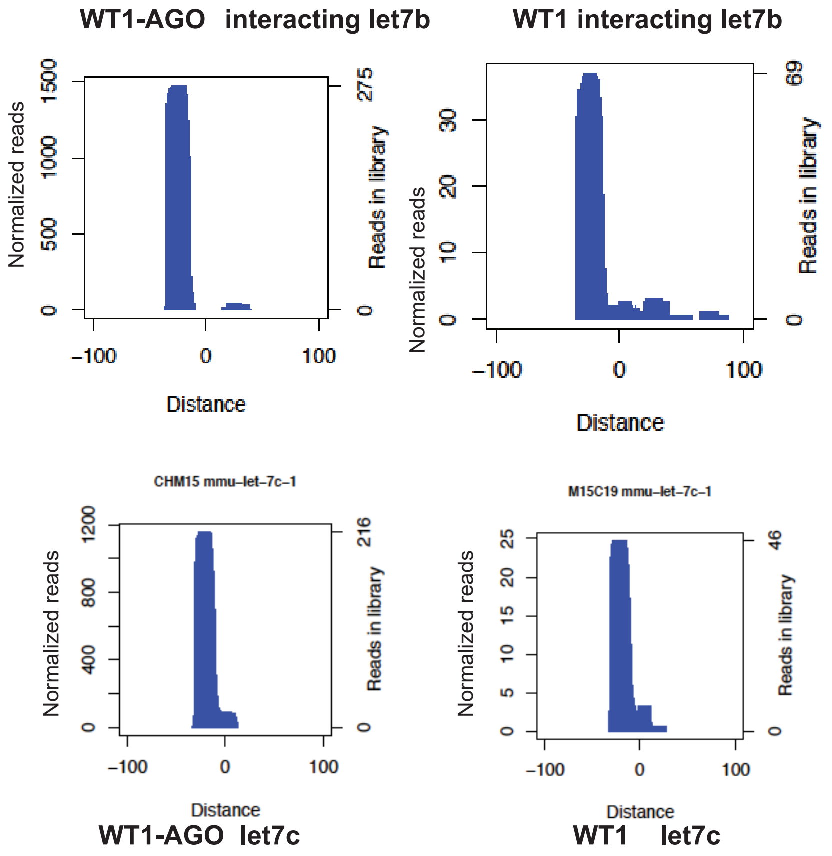
WT1-AGO interacting let7b
WT1 interacting let7b
Normalized reads
Reads in library
Distance
−100
0
100
1500
1000
500
275
30
20
10
69
CHM15 mmu–let–7c–1
M15C19 mmu–let–7c–1
1200
800
400
216
25
20
15
10
5
46
WT1-AGO let7c
WT1 let7c

**FIGURE S4**

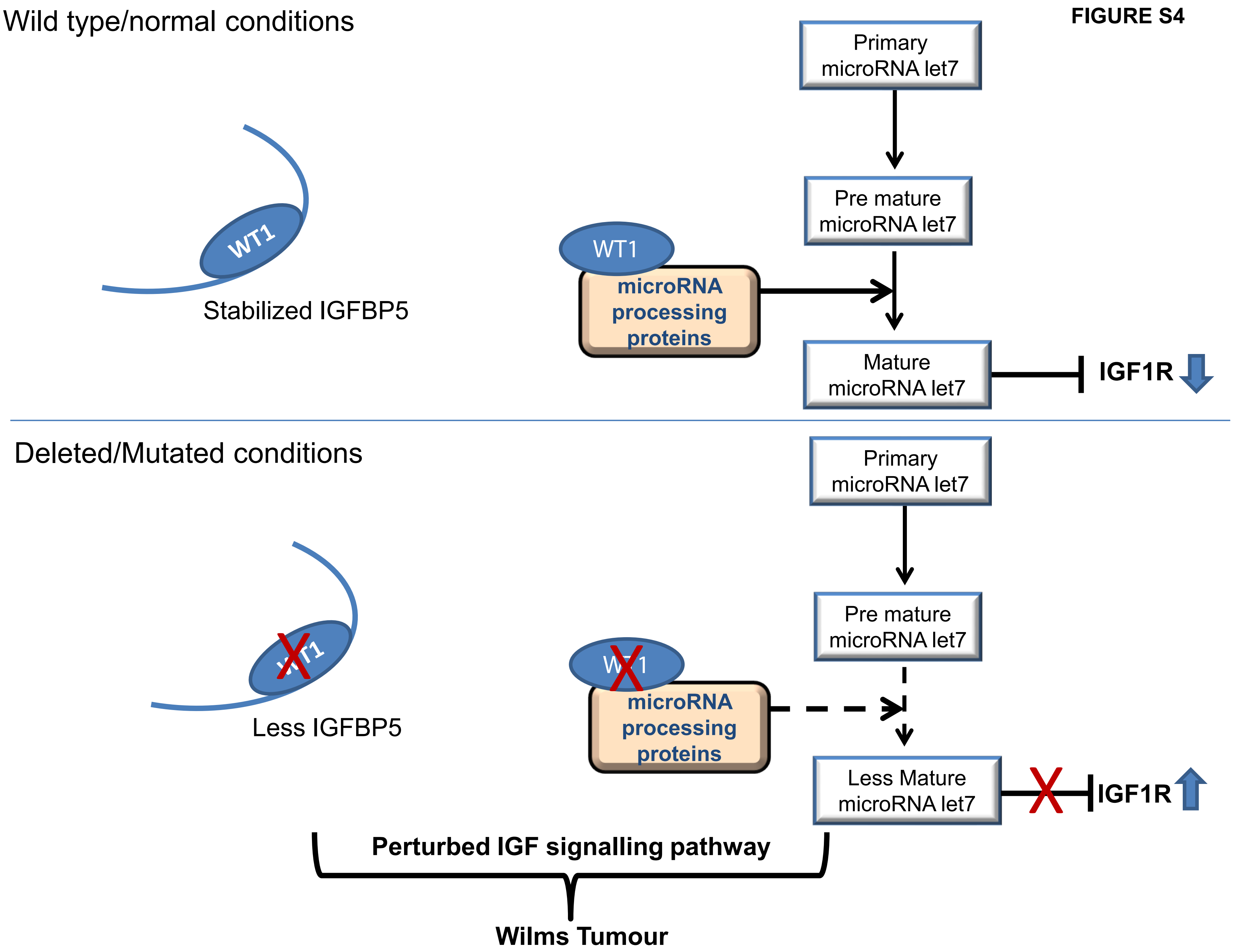
Wild type/normal conditions
Primary microRNA let7
Pre mature microRNA let7
WT1
Stabilized IGFBP5
WT1
microRNA processing proteins
Mature microRNA let7
IGF1R
Deleted/Mutated conditions
Primary microRNA let7
Pre mature microRNA let7
Less IGFBP5
microRNA processing proteins
Less Mature microRNA let7
IGF1R
Perturbed IGF signalling pathway
Wilms Tumour

Table S1

| miRNA-mRNA hybrids from FLASH | | | mRNA fold change from transcriptome analysis | |
|---|---|---|---|---|
| miRNA | protein-codingGene | energy | log2Fold Change | p_adj |
| Gm25732 | Ankrd1 | -7.5 | 3.76956 | 0.029538 |
| Gm24271 | D0H4S114 | -8.1 | -4.33955 | 0.0197962 |
| Mir199a-2 | Neto1 | -11 | -4.85571 | 0.120975 |
| Gm27537 | Serinc5 | -4 | -3.32667 | 0.0961279 |
| Mir26a-2 | Cfh | -15 | -6.25808 | 0.0004408 2 |
| Mir21a | Cdh11 | -12.6 | -4.73714 | 0.0041257 1 |
| Mir351 | Btg2 | -5.6 | -3.51285 | 0.105283 |
| Mir351 | Ypel2 | -10.2 | -3.33725 | 0.112054 |

**Table S2**

**Oligo details:**

**a) Oligos for northern probes (5’-3’)**

| | |
|---|---|
| **let7c** | TGAGGTAGTAGGTTGTATGGTTCCTGTCTC |
| **let7g** | TGAGGTAGTAGTTTGTACAGTTCCTGTCTC |
| **U6** | GCTAATCTTCTCTGTATCGTTCCAATTTTAGTATATGTGCTGCCG |

**b) Gene specific primers**

| | FP (5’-3’) | RP (5’-3’) |
|---|---|---|
| **Wt1** | GCCTTCACCTTGCACTTCTC | GACCGTGCTGTATCCTTGGT |
| **Fgf5** | AGTCAATGGCTCCCACGAAG | GGCACTTGCATGGAGTTTTCC |
| **Igf1r** | TTTCCACTCCGCATTTCTGC | CGTCCAAAAACAAGAGCGCA |
| **Hmga2** | ACCTGTGAGCCCTCTCCTAA | CACCTGGTGCGTTCTCCAAA |
| **Lin28a** | CTTTGTGCACCAGAGCAAGC | CGCAGTTGTAGCACCTGTCT |
| **Lin28b** | ACTCCGGTCCAAAGGGAAAG | GGCTCTTCACCTTTGCTTGC |
| **Pou5f1** | AAGTTGGCGTGGAGACTTTG | TTCATGTCCTGGGACTCCTC |
| **CCR7** | CATGGACCCAGGTGTGCTTC | TCAGTATCACCAGCCCGTTG |
| **Col1a2** | CACCCCAGCGAAGAACTCAT | TCTCCTCATCCAGGTACGCA |
| **Yy1** | ACAGGCAAGAAACTCCCTCC | CTTTGAGCTCTCAACGAACGC |
| **Fgf11** | CCAGCTCCTTCACCCACTTC | ACGCACTCCTTAAAGCGACA |
| **Hnrnpa1** | ACGGAACCAAGGTGGCTATG | TCCCTGTCACTTCTCTGGCT |
| **Tet2** | TGAATATTGATGCGGAGGCGA | CTCAGCATGGGTGGTTCTGT |
| **Igf2** | CCCCAGCCCTAAGATACCCTAA | GTATGCAAACCGAACAGCGG |
| **Mef2c** | GCAGAGGGATCACGCATCTC | TGCCTTTCTGCTTCTCCAGG |
| **Dzip** | TGCGACTGTGGCTGCTTATTA | TGGCCTGACTCTCGATTCCT |
| **Robo1** | GGCAGTGACTGTGGATGACA | TTACAACGAAATGTGGCGGC |
| **Wnt1** | TGATGTTTGCCCACCCTACC | CCTCAGGATGGCAAAAGGGT |
| **Tug1** | AGTGTACTCTGCTGTCAACCT | GTTTCAGCCTCTCCTTTGCC |
| **Birc5** | CGAACTCAGAAATCCGCCTG | AGCTCAATTCCAGGGACCAA |
| **S14** | ATTGCAGGCTAGAGTTGGGT | GCCTCACAGCCTGTCTAGAA |
| **Malat1** | GGTGGGCTTTTGTTGATGAG | CAAACGAAACATTGGCACAC |
| **Neat1** | GACAACTGCGGTGTTCTGTT | AAGTAGTCTCCATGGGCCAC |
| **18s** | GTAACCCGTTGAACCCCATT | CCATCCAATCGGTAGTAGCG |
| **GAPDH** | ATGTAGGCCATGAGGTCCAC | GGGATGGGAAACCTGACTTT |

**c) miRNA primers:**

| | | |
|---|---|---|
| **pre let7c** | TGTGTGCATCCGGGTTG | AGTGTGCTCCAAGGAAAG |
| | GCATCCGGGTTGAGGTAGTAG | GCTCCAAGGAAAGCTAGAAG |
| **pre let7g** | CAGTAAGAAATATGGTGTGGACCTCA | GATGTATTTGTTGTCATGCAAGGAC |
| **pre miR 125b** | TGCGCTCCCCTCAGTCCCTGAGAC | CAGCTCCCAAGAGCCTAACCCGTG |

**let7c** TGAGGTAGTAGGTTGTATGGTT

**let7g** TGAGGTAGTAGTTTGTACAGTT

**miR21** TAGCTTATCAGACTGATGTTGA

**miR26** TTCAAGTAATCCAGGATAGGCT

**miR92** AGGTTGGGATTTGTCGCAATGCT

**miR99** AACCCGTAGATCCGATCTTGTG

**miR125b**TCCCTGAGACCCTAACTTGTGA

**miR101** TCAGTTATCACAGTGCTGATGC

**miR181** AACATTCAACGCTGTCGGTGAGT
